# Recessive inactivating mutations in *TBCK*, encoding a Rab GTPase-activating protein that modulates mTOR signaling, cause severe infantile syndromic encephalopathy

**DOI:** 10.1101/036111

**Authors:** Jessica X. Chong, Viviana Caputo, Ian G. Phelps, Lorenzo Stella, Lisa Worgan, Jennifer C. Dempsey, Alina Nguyen, Vincenzo Leuzzi, Richard Webster, Antonio Pizzuti, Colby T. Marvin, Gisele E. Ishak, Simone Ardern–Holmes, Zara Richmond, Univ of Washington Center for Mendelian Genomics, Michael J. Bamshad, Xilma R. Ortiz-Gonzalez, Marco Tartaglia, Maya Chopra, Dan Doherty

## Abstract

Infantile encephalopathies are a group of clinically and biologically heterogeneous disorders for which the genetic basis remains largely unknown. Here, we report a previously unrecognized syndromic neonatal encephalopathy characterized by profound developmental disability, severe hypotonia, seizures, diminished respiratory drive requiring mechanical ventilation, brain atrophy, corpus callosum dysgenesis, cerebellar vermis hypoplasia, and facial dysmorphism. Biallelic inactivating mutations in *TBCK* (TBC1 domain-containing kinase) were independently identified by Whole-Exome Sequencing (WES) as the cause of this condition in four unrelated families. Matching these families was facilitated by sharing phenotypic profiles and WES data in a recently released web-based tool (Geno_2_MP) that links phenotypic information to rare variants in families with Mendelian traits. TBCK is a putative GTPase-activating protein (GAP) for small GTPases of the Rab family and has been shown to control cell growth and proliferation, actin cytoskeleton dynamics, and mTOR signaling. Two of the three mutations are predicted to truncate the protein (c.376C>T [p.Arg126*] and c.1363A>T [p.Lys455*]), and loss of the major TBCK isoform was confirmed in primary fibroblasts from one affected individual. The third mutation, c.1532G>A [p.Arg511His], alters a conserved residue within the TBC1 domain. Structural analysis implicates Arg^511^ as a required residue for Rab-GAP function, and *in silico* homology modeling predicts impaired GAP function in the corresponding mutant. These results suggest loss of Rab-GAP activity is the underlying mechanism of disease. In contrast to other disorders caused by dysregulated mTOR signaling associated with focal or global brain overgrowth, impaired TBCK function results in progressive loss of brain volume.

---

Severe infantile encephalopathy is a non-specific clinical condition that can be caused by hypoxemia, hemorrhage, toxins (e.g., hyperbilirubinemia or withdrawal from selective serotonin reuptake inhibitors or narcotics), and mutations in multiple genes such as *PURA* [MIM 600473], *UNC80* [MIM 612636] or *NALCN* [MIM 611549].^1–9^ Over the past 5 years, substantial progress has been made toward defining the genetic basis of infantile encephalopathies; however, the genetic cause remains unknown for a large proportion of affected individuals, and these conditions have few distinguishing clinical features, making it challenging to stratify affected individuals for gene discovery efforts. Whole-Exome Sequencing (WES) of multiple independent families with non-specific infantile encephalopathies is a powerful way to discover shared candidate genes among affected individuals, which upon further clinical evaluation are often found to share phenotypic features delineating a distinctive condition within this otherwise non-specific category. Here, we report on a previously unrecognized syndromic infantile encephalopathy characterized by severe developmental disability, brain atrophy, focal seizures with early-onset, central respiratory failure, and facial dysmorphism. WES in four families led to the discovery of mutations in *TBCK*, a gene encoding a putative Rab-specific GTPase-activating protein (GAP), as the underlying molecular cause.

Clinical data and biological samples were obtained from three unrelated families after written informed consent was provided, and studies for each family were approved by the respective institutional review boards of the University of Washington (Seattle, USA), the Genetics Services Advisory Committee (New South Wales, Australia), and Università La Sapienza (Rome, Italy). A single affected individual with neonatal onset encephalopathy, brain atrophy with cerebellar malformation, and later-onset seizures (Figure 1, Table 1, and Table S2 Family A) was referred to one of us with the tentative diagnosis of Joubert syndrome [MIM PS213300] with atypical features. Twenty-seven genes previously implicated in Joubert syndrome were screened,^10^ but no predicted-pathogenic mutations were found. Upon re-evaluation, the clinical and MRI and features were inconsistent with the diagnosis of Joubert syndrome (i.e., early respiratory failure, epilepsy, developmental regression, dysmorphic features, and absence of classic imaging features), which suggested that this person instead had a condition involving hindbrain malformation that had not been previously delineated. Subsequently WES was performed by the University of Washington Center for Mendelian Genomics (UW-CMG) as described previously.^11^ Single nucleotide variants (SNVs) were annotated with the SeattleSeq137 Annotation Server. SNVs categorized as intergenic, coding-synonymous, UTR, near-gene, or intron were excluded. Variants flagged as low-quality or potential false positives (quality score ≤ 50, long homopolymer run > 4, low quality by depth < 5, within a cluster of SNPs), as well as those with a mean alternative allele frequency >0.005 in the NHLBI Exome Sequencing Project Exome Variant Server (ESP6500/EVS) or in an internal WES database of ~700 exomes, were also excluded. Allowing for recessive and X-linked inheritance, this filtering left 30 candidate genes (Table S1); however, 27 genes could be excluded based on one of the variants being present homozygous in the ExAC database or having a CADD score <15.^12^ Only one gene, *TBCK*, harbored a biallelic truncating variant (hg19 chr4:107183260 G>A; RefSeq: NM_001163435.2 c.376C>T, p.Arg126*), so *TBCK* was considered the best candidate. No other individuals with *TBCK* mutations were known at the time, so this candidate gene was archived in 2013 internally and later in Geno_2_MP,^13^ a recently-released web-based tool/database developed by the UW-CMG that links phenotypic descriptions to rare-variant genotypes in ~3,800 exomes from families with Mendelian conditions.

**Figure 1.**
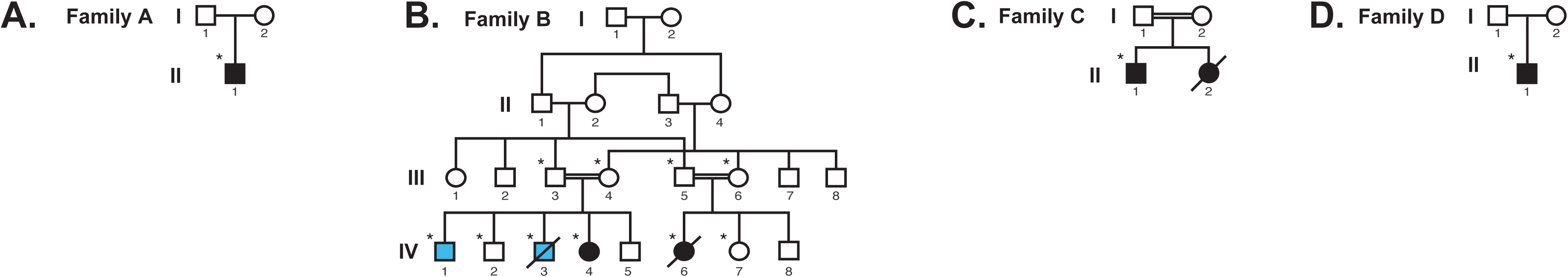
Pedigrees and pictures of individuals with *TBCK-related* encephalopathy (photographs removed at request of bioRxiv). (**A-D**) Pedigrees for four families segregating loss of function mutations in *TBCK*. Solid black fill indicates individuals affected with TBCK-related encephalopathy. Light blue fill in Family B indicates individuals affected by a blood-related phenotype without neurodevelopmental features; neither was homozygous for a mutation in *TBCK*. WES was performed on individuals marked with *’s. (E-J) Similar facial features and hypotonia in Family A-II-1 at 25 months **(E)** and 13 years **(F)**, Family B-IV-4 at 17 months **(G)** and 4 years 3 months **(H)**, Family B-IV-6 at 18 months **(I)**, Family C-II-1 at 21 months **(J)**, and Family D-II-1 at 14 years. See Table 1 for detailed clinical information for each affected individual and Figure 2 and Table S3 for imaging information.

**Table 1.**
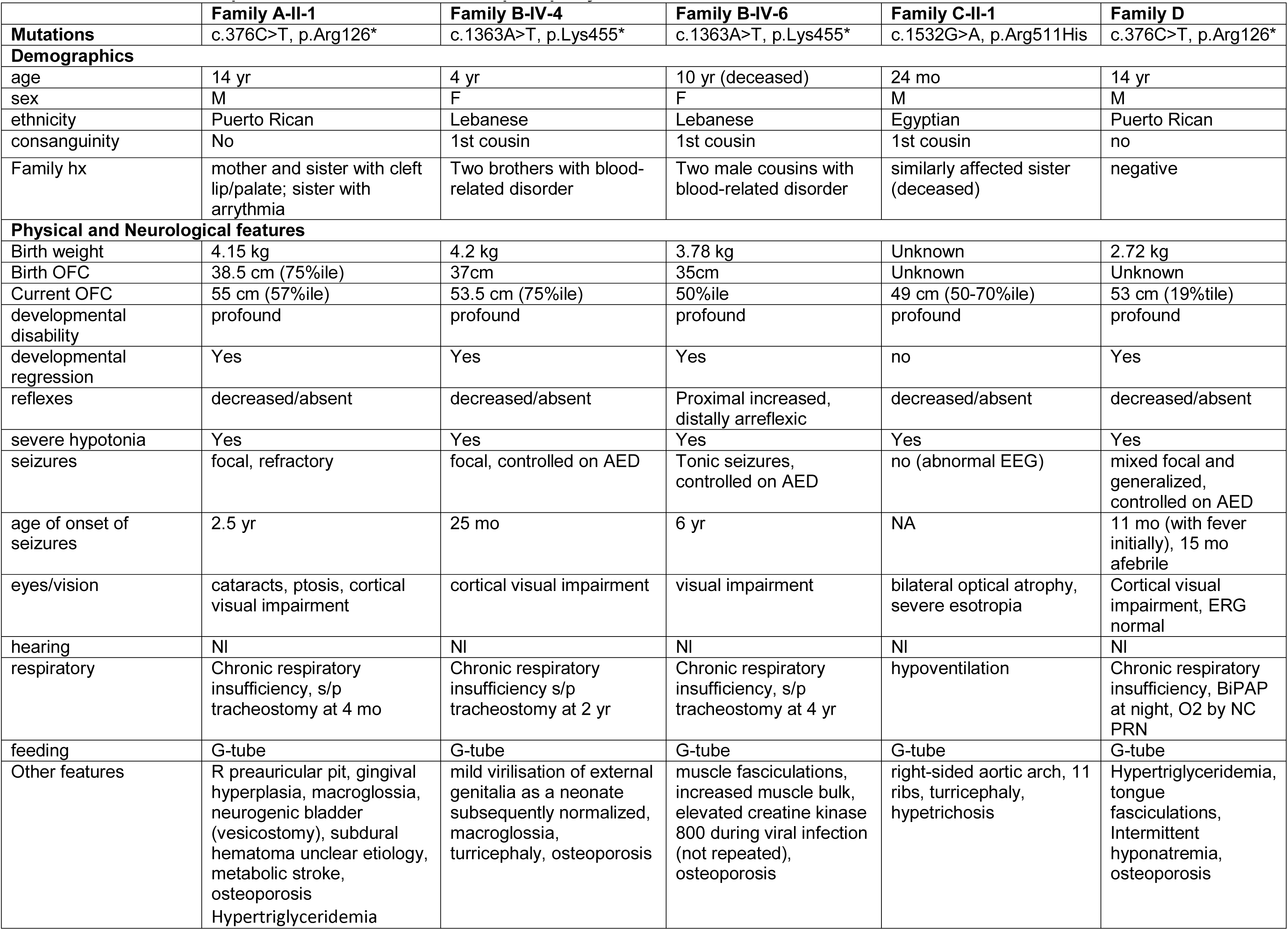
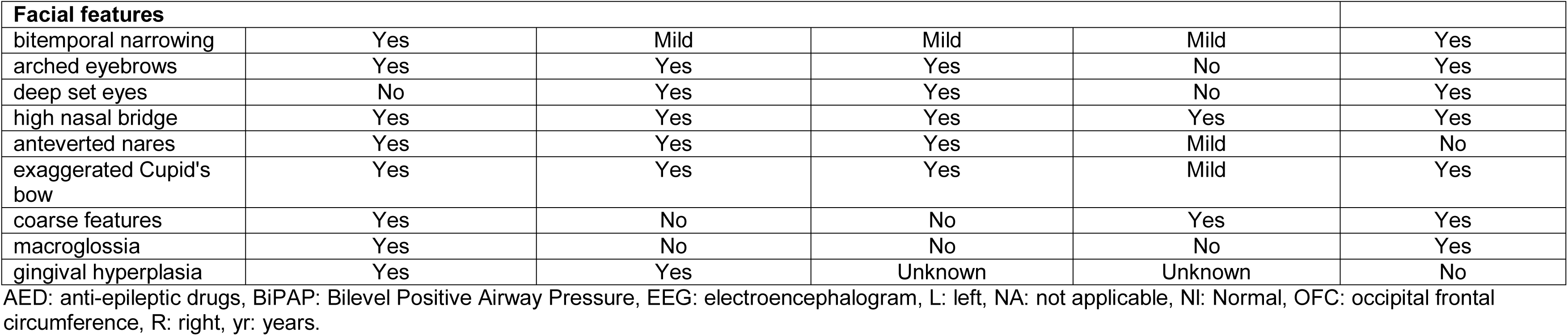
Clinical features in patients with *TBCK*-related encephalopathy

|  | **Family A-II-1** | **Family B-IV-4** | **Family B-IV-6** | **Family C-II-1** | **Family D** |
| --- | --- | --- | --- | --- | --- |
| **Mutations** | c.376C>T, p.Arg126* | c.1363A>T, p.Lys455* | c.1363A>T, p.Lys455* | c.1532G>A, p.Arg511His | c.376C>T, p.Arg126* |
| **Demographics** |  |  |  |  |  |
| age | 14 yr | 4 yr | 10 yr (deceased) | 24 mo | 14 yr |
| sex | M | F | F | M | M |
| ethnicity | Puerto Rican | Lebanese | Lebanese | Egyptian | Puerto Rican |
| consanguinity | No | 1st cousin | 1st cousin | 1st cousin | no |
| Family hx | mother and sister with cleft lip/palate; sister with arrhythmia | Two brothers with blood-related disorder | Two male cousins with blood-related disorder | similarly affected sister (deceased) | negative |
| **Physical and Neurological features** |  |  |  |  |  |
| Birth weight | 4.15 kg | 4.2 kg | 3.78 kg | Unknown | 2.72 kg |
| Birth OFC | 38.5 cm (75%ile) | 37cm | 35cm | Unknown | Unknown |
| Current OFC | 55 cm (57%ile) | 53.5 cm (75%ile) | 50%ile | 49 cm (50-70%ile) | 53 cm (19%tile) |
| developmental disability | profound | profound | profound | profound | profound |
| developmental regression | Yes | Yes | Yes | no | Yes |
| reflexes | decreased/absent | decreased/absent | Proximal increased, distally areflexic | decreased/absent | decreased/absent |
| severe hypotonia | Yes | Yes | Yes | Yes | Yes |
| seizures | focal, refractory | focal, controlled on AED | Tonic seizures, controlled on AED | no (abnormal EEG) | mixed focal and generalized, controlled on AED |
| age of onset of seizures | 2.5 yr | 25 mo | 6 yr | NA | 11 mo (with fever initially), 15 mo afebrile |
| eyes/vision | cataracts, ptosis, cortical visual impairment | cortical visual impairment | visual impairment | bilateral optical atrophy, severe esotropia | Cortical visual impairment, ERG normal |
| hearing | NI | NI | NI | NI | NI |
| respiratory | Chronic respiratory insufficiency, s/p tracheostomy at 4 mo | Chronic respiratory insufficiency s/p tracheostomy at 2 yr | Chronic respiratory insufficiency, s/p tracheostomy at 4 yr | hypoventilation | Chronic respiratory insufficiency, BiPAP at night, O2 by NC PRN |
| feeding | G-tube | G-tube | G-tube | G-tube | G-tube |
| Other features | R preauricular pit, gingival hyperplasia, macroglossia, neurogenic bladder (vesicostomy), subdural hematoma unclear etiology, metabolic stroke, osteoporosis<br>Hypertriglyceridemia | mild virilisation of external genitalia as a neonate subsequently normalized, macroglossia, turricephaly, osteoporosis | muscle fasciculations, increased muscle bulk, elevated creatine kinase 800 during viral infection (not repeated), osteoporosis | right-sided aortic arch, 11 ribs, turricephaly, hypetrichosis | Hypertriglyceridemia, tongue fasciculations, Intermittent hyponatremia, osteoporosis |

| Facial features |  |  |  |  |  |
| --- | --- | --- | --- | --- | --- |
| bitemporal narrowing | Yes | Mild | Mild | Mild | Yes |
| arched eyebrows | Yes | Yes | Yes | No | Yes |
| deep set eyes | No | Yes | Yes | No | Yes |
| high nasal bridge | Yes | Yes | Yes | Yes | Yes |
| anteverted nares | Yes | Yes | Yes | Mild | No |
| exaggerated Cupid's bow | Yes | Yes | Yes | Mild | Yes |
| coarse features | Yes | No | No | Yes | Yes |
| macroglossia | Yes | No | No | No | Yes |
| gingival hyperplasia | Yes | Yes | Unknown | Unknown | No |

AED: anti-epileptic drugs, BiPAP: Bilevel Positive Airway Pressure, EEG: electroencephalogram, L: left, NA: not applicable, NI: Normal, OFC: occipital frontal circumference, R: right, yr: years.

In late 2014, two affected cousins in a second family were ascertained on the basis of an apparently-novel neurodegenerative phenotype including cerebellar hypoplasia (Figure 1, Table 1, Table S2, Family B). Samples from the extended family underwent high-density genotyping and WES by the UW-CMG as described previously.^11^ Linkage analysis conducted under a recessive model, assuming a causal allele frequency of 0.0001 and full penetrance (f=0,0,1.0), yielded two peaks (hg19: chr4:88837705-109066366 and chr7:81735015-112087033) with a maximum parametric LOD score of 2.48. WES data were annotated with the SeattleSeq138 Annotation Server. As with the analysis for Family A, variants unlikely to impact protein coding sequence, variants flagged by GATK as low quality, and variants with an alternative allele frequency >0.002 in any population in the ESP6500/EVS, 1000 Genomes (phase 3 release), or Exome Aggregation Consortium (ExAC v1.0) browser were excluded. Copy number variant (CNV) calls were generated from exome data with CoNIFER.^14^ After overlapping the linkage and exome data, only three genes, *VPS50* [MIM 616465] (hg19 chr7:92953034 G>A; Refseq: NM_001257998.1, c.1877G>A, p.Arg626Gln), *LRCH4* (hg19 chr7:100173865 C>T; Refseq: NM_002319.3, c.1634G>A, p.Arg545His; rs370008127), and *TBCK* (hg19 chr4:107156512 T>A; RefSeq: NM_001163435.2, c.1363A>T, p.Lys455*; rs376699648), remained as candidates. All three genes were submitted to GeneMatcher and Geno_2_MP. No matches were found for *VPS50* or *LRCH4* in either database, but Geno_2_MP yielded a single individual (Figure 1 and Table 1, Family A) with “cerebellar malformation” who was homozygous for a nonsense variant in *TBCK*. Sharing of the same candidate gene and an overlapping phenotype suggested that the putative loss of function mutations in *TBCK* were causal.

In parallel, a third family was ascertained due to profound developmental disability associated with brain atrophy before two years of age in a son, and death of a 12 month old daughter exhibiting similar features (Figure 1, Table 1, and Table S2, Family C). WES was performed on a sample from the affected son (Family C-II-1). Variants were called using an in-house pipeline,^15–17^ and filtered against public and in-house databases, retaining only clinically-associated variants and variants with MAF <0.001 (dbSNP142), <0.002 (ExAC), <0.01 (in-house database, ~500 exomes), or unknown MAF. The SnpEff toolbox (v4.1) was used to predict the impact of variants, which were filtered to retain only functionally relevant variants *(i.e*., missense, nonsense, coding indel variants, and intronic variants located from −5 to +5 with respect to an exon-intron junction). Functional annotation of variants was performed using SnpEff v4.1 and dbNSFP2.8. Parental first cousin consanguinity and the death of a female sibling with similar clinical features *(i.e*., suggestive facies, severe developmental delay, generalized hypotonia and congenital heart malformation) strongly suggested autosomal recessive inheritance, and based on the hypothesis of homozygosity by descent, 16 candidate genes were identified. These candidates were stratified through a mixed filtering/prioritization strategy designed to retain genes with predicted-damaging variants (CADD_phred > 15), ranked on the basis of their biological relevance on the developmental processes altered in the disorder (GeneDistiller) (see Table S1).^18^ Among the 8 retained genes, *TBCK* (hg19 chr4:107154202 C>T; RefSeq: NM_001163435.2 c.1532G>A, p.Arg511His) received the highest score from GeneDistiller and was considered the best candidate. At this point, M.T. contacted M.J.B. and inquired whether *TBCK* was a candidate gene for any phenotypes being studied at the UW-CMG.

Another individual of Puerto Rican descent (Figures 1 and 2, Family D II-1) with severe hypotonia, chronic respiratory insufficiency requiring nighttime BiPAP (Bilevel Positive Airway Pressure), brain atrophy, and similar facial features (Family D-II-1 in Figure 1) was also ascertained in parallel. Clinical exome sequencing identified the same homozygous p.Arg126* variant present in Family A-II-1, supporting the existence of a founder mutation in the Puerto Rican population. The phenotypic features in Family D-II-1 are quite similar to the other affected individuals (Tables 1, S2, and S3).

**Figure 2.**
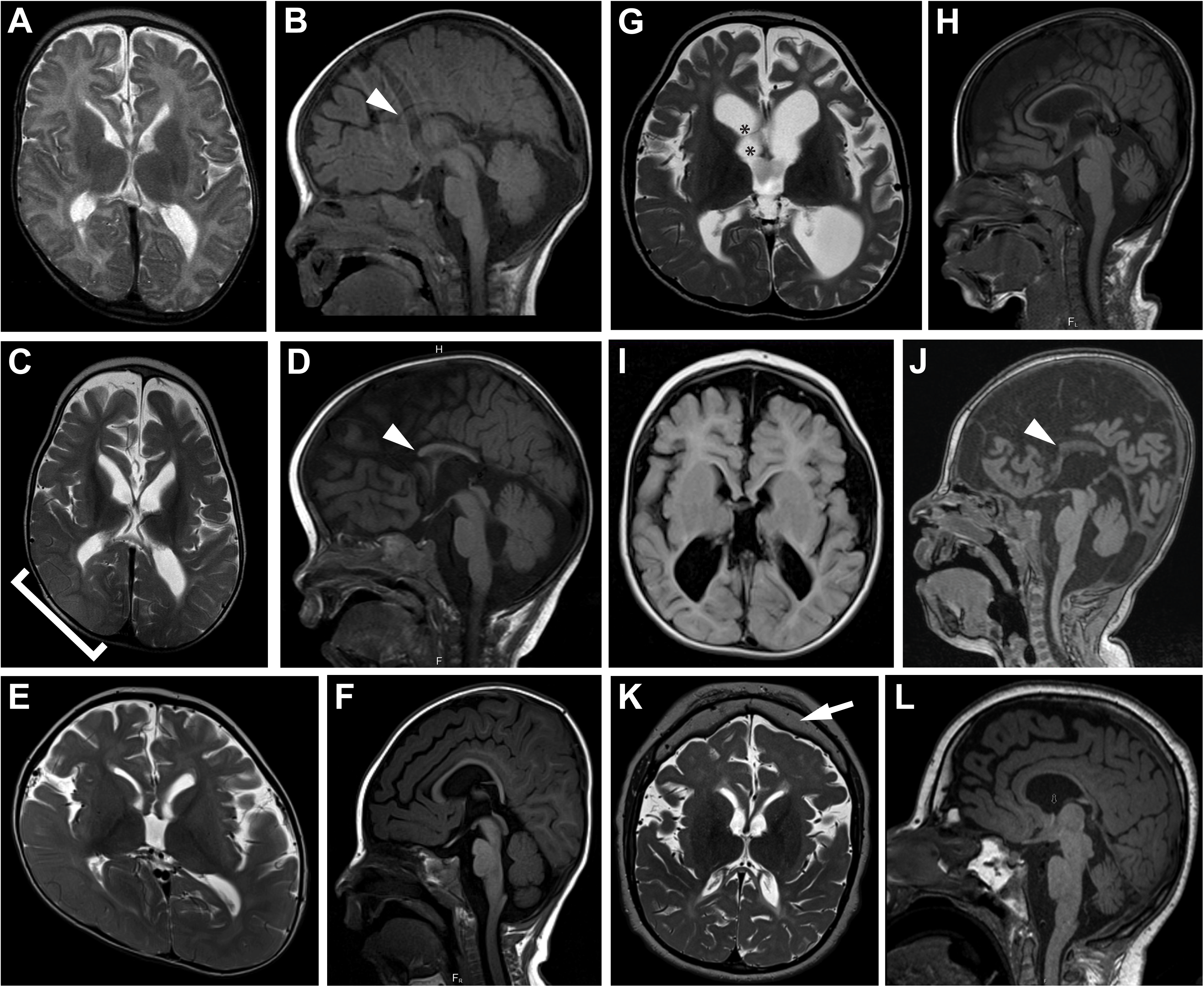
Brain imaging features in individuals with TBCK-related encephalopathy. **(A-D)** Progressive gray and white matter volume loss, most severe in the frontal lobes, demonstrated by increasing ventriculomegaly and extraxial spaces (cortical and cerebellar) in Family A-II-1 between 22 days of age **(A-B)** and 29 months of age **(C-D)**. Swelling of the right parieto-occipital lobe, presumably due to a metabolic stroke (bracket in C), diffusely thin corpus callosum with absent rostrum (arrowhead in B and D), and mild cerebellar vermis hypoplasia with relatively spared brainstem are also present. **(E-F)** Ventriculomegaly, prominent extraxial spaces, diffusely thin but complete corpus callosum, and mild cerebellar vermis hypoplasia in Family B-IV-4 at 18 months of age. Right plagiocephaly is also present. **(G-H)** Marked ventriculomegaly, prominent extraxial spaces, diffusely thin but complete corpus callosum, and mild cerebellar vermis hypoplasia in Family B-IV-6 at 6 years of age. Synechiae are also present in the right frontal horn (asterisks in G). **(I-J)** Ventriculomegaly, prominent extraxial spaces, diffusely thin corpus callosum with absent rostrum and anterior body (arrowhead in J), and mild cerebellar vermis hypoplasia in Family C-II-1 at 21 months of age. **(K-L)** Mild ventriculomegaly, prominent extraxial spaces, diffusely thin corpus callosum, mild cerebellar vermis hypoplasia with relatively preserved brainstem, and thick frontal bone (arrow in K) in Family D-II-1 at 14 years of age. (**A, C, E, G, K**) axial T2-weighted images, **(I)** axial T1-weighted image, (**B, D, F, H, J, L**) sagittal T1-weighted images.

The variants in each family were either rare or not present in reference population databases. Specifically, the maximum alternate allele frequencies in any population from EVS, 1000 Genomes phase 3, or ExAC v0.3 was 0.5208% for p.Arg126* (Families A and D) and 0.0107% for p.Lys455* (Family B) in ExAC Latinos, while p.Arg511His (Family C) was not present in any database. Each variant had a high CADD score:^12^ 37.0 for p.Arg126*, 42.0 for p.Lys455*, and 35.0 for p.Arg511His.

All affected individuals displayed profound developmental disability, making little progress beyond infancy, and a relatively homogeneous phenotype (Table 1, Figure 1). Additionally, Family A-II-1 and Family B-IV-4 and IV-6 exhibited developmental regression (loss of visual fixation and following). All four individuals also required gastrostomy tube feedings and had decreased respiratory drive, with the three older children requiring tracheostomies for chronic mechanical ventilation. Family A-II-1, Family B-IV-4 and Family B-IV-6 also developed focal seizures between 2 and 6 years of age, and Family C-II-1 had an abnormal electroencephalogram before 2 years of age without clinical seizures. Family A-II-1, Family B-IV-4 and Family C-II-1 had decreased to absent reflexes, while Family B-IV-6 developed hyperreflexia. Shared facial features included bitemporal narrowing, arched eyebrows, deep set eyes, high nasal bridge, anteverted nares, and an exaggerated “Cupid's bow” of the upper lip (Figure 1D-H). Family A-II-1 and Family B-IV-6 developed osteoporosis, which could be either a non-specific effect of chronic, severe disability or a more specific feature of *TBCK*-related disease. At three years of age, Family A-II-1 had an acute neurological decompensation with restricted diffusion of the right parieto-occipital cortex, indicative of a metabolic stroke, from which he recovered to his baseline neurological status. In all four individuals, brain imaging revealed increased ventricular and extra-axial spaces and diffusely decreased white matter volume, with atrophy confirmed by serial MRI in Family A-II-1 and Family B-IV-4 (Figure 2 and Table S3). Despite the loss of brain volume, microcephaly was not observed. In addition, all individuals have corpus callosum dysgenesis (thinning, partial agenesis or both), increased T2/FLAIR signal in the periventricular white matter, and cerebellar vermis hypoplasia.

*TBCK* encodes a protein with a TBC (Tre-2, Bub2, and Cdc16) domain flanked by an *N*-terminal kinase-like domain and a rhodanese homology domain at the *C*-terminus, but its function has not been extensively characterized. Due to the lack of key catalytic residues, the kinase domain is presumably inactive.^19^ Multiple *TBCK* mRNAs have been reported with two major isoforms differing in the presence or absence of the *N*-terminal portion that encodes the kinase-like domain (Figure 3A).^20^ To characterize the impact of the truncating mutations, the TBCK protein level was evaluated by western blot using skin fibroblasts obtained from an individual who was homozygous for p.Arg126*, Family A-II-1 (Figure 3B). As expected, two major bands at ~101 and 71 kD, representing respectively the long (full-length) protein, and the shorter isoform without the *N*-terminal kinase-like domain,^20^ were observed in two control fibroblast lines, with the full-length isoform representing the more abundant product. In contrast, the full-length protein was nearly absent in the fibroblasts from Family A-II-1, and the 71 kD isoform was also reduced. While the transcript for the 71kD isoform is not directly affected by the mutation, the reduced protein level indicates that the mutation perturbs the levels of both major *TBCK* isoforms in fibroblasts.

**Figure 3.**
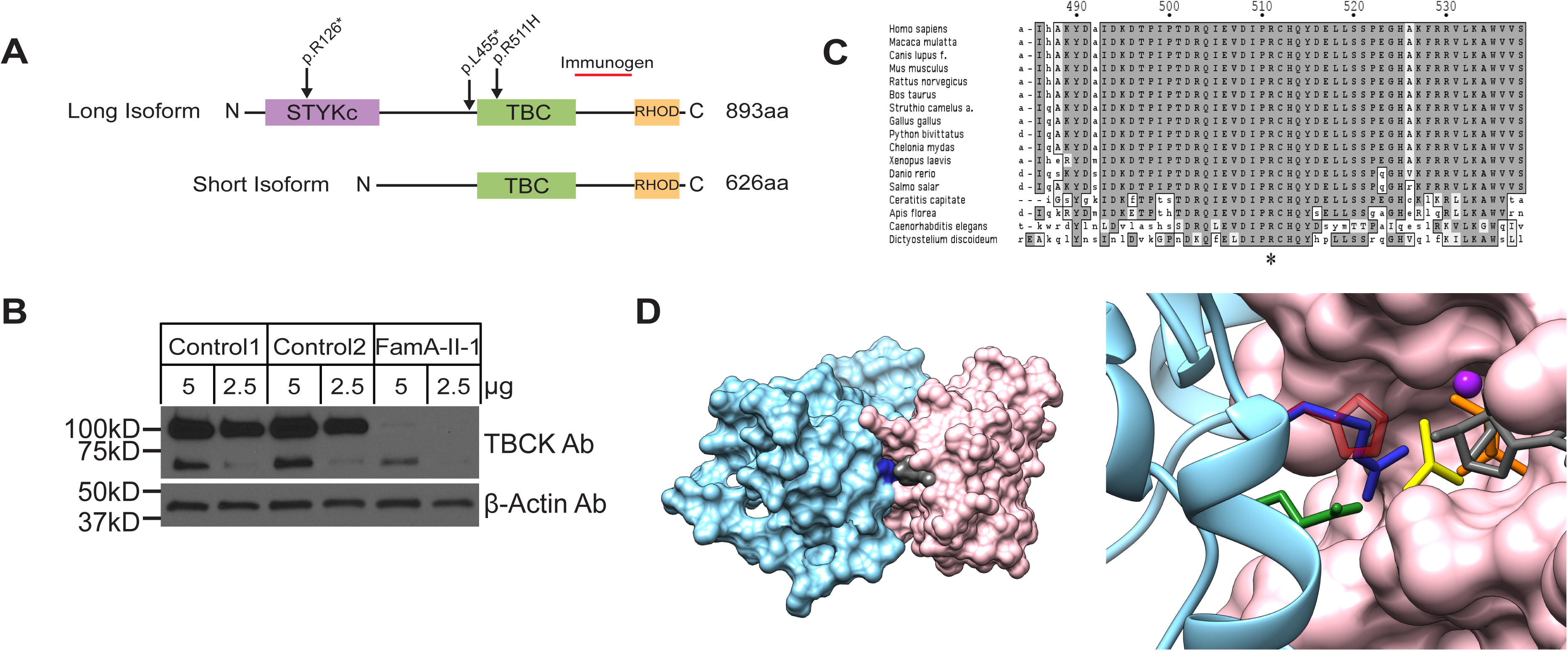
Biochemical and structural characterization of *TBCK* mutations. **(A)** *TBCK* encodes two isoforms containing a TBC1 domain (TBC) flanked by a rhodanese domain (RHOD) at the *C*-terminus. The long isoform also contains a pseudokinase domain (STYKc) at the *N*-terminus. Location of the identified mutations is reported. **(B)** Western blot showing dramatically reduced levels of TBCK in cell line from Family A-II-1 versus two control cell lines. The TBCK monoclonal antibody (1/250, Sigma HPA039951) recognizes a *C*-terminal fragment of the protein depicted as a red line in **(A)**. Following stripping, (3-actin (1/5,000, Sigma A5441) was used as a loading control. Predicted sizes of TBCK long and short isoforms are 101kD and 71kD respectively. **(C)** Conservation of the catalytic arginine “finger” (Arg^511^ in TBCK) in TBCK orthologs. **(D)** Homology model of the TBCK TBC1 domain complexed with the Rab33 GTPase. In the overall structure of the complex (left), both proteins are shown in a surface representation, with Rab colored in pink, GDP in gray, the TBC1 domain in light blue and Arg^511^ in blue. In the enlarged view of the active site of the GTPase (right), GDP is reported in gray sticks, with the phosphates colored in orange. AlF_3_, which in the crystal structure mimics the transition state for GTP hydrolysis, is shown in yellow and the Mg atom as a purple sphere. The TBC1 domain is reported in ribbon representation, with the side-chains of key residues shown as sticks. The two catalytic “fingers” Arg^511^ (blue) and Gln^546^ (green) are shown together with the disease-associated His^511^ reported in semitransparent red.

The c.1532G>A mutation in Family C-II-1 is predicted to alter a highly conserved arginine residue located in the TBC1 domain (Figure 3C). This structural unit of approximately 200 amino acids is characteristic of most GAPs that regulate the Rab family of small GTPases.^21,22^ TBC domains negatively control Rab function by promoting GTP hydrolysis *via* stabilization of the transition state of the reaction. Specifically, the TBC domain interacts with the substrate, and catalyzes the reaction by using a so-called “dual finger” mechanism, in which a key Arg residue projects into the Rab active site.^23–25^ Since no crystallographic structure of TBCK was available, a homology model of the TBC1 domain of TBCK (residues 467–648) in complex with a Rab protein was constructed to explore the structural impact of the disease-causing amino acid substitution. This model was generated using DeepView and the SwissModel server,^26^ based on the crystallographic structure of the Gyp1 TBC domain (36% sequence identity with the TBCK TBC domain) complexed with Rab33 bound to GDP and AlF**3** (PDB code 2G77).^23^ TBCK was originally incorrectly classified as an unconventional TBC protein lacking the “arginine finger”;^21^ however, our modeling data support more recent bioinformatics analyses^21^ indicating that Arg^511^ does in fact represent an arginine finger essential for GAP function. The p.Arg511His substitution was introduced *in silico* with UCSF Chimera, ^27^ and even though the Arg-to-His substitution introduces a side chain that could maintain the overall positive charge under appropriate conditions,^28^ the introduced residue is significantly bulkier and shorter than Arg, and unable to project into the GTPase active site. In the model, the minimum distance between Arg^511^ and the GDP phosphate is 3 Å, while this distance increases to 7 Å in the mutant. These considerations strongly point to impaired GAP activity as the predicted mechanism of disease associated with the p.Arg511His substitution.

TBCK was recently documented to play a role in the control of cell proliferation, cell size, and actin cytoskeleton dynamics.^19,20^ While the lack of several residues with critical function in mediating the activity of the kinase domain supports the view that TBCK is catalytically inactive,^29,30^ the conservation of the key catalytic residues of the TBC1 domain, as well as the identification of a disease-causing mutation specifically targeting one of the two key-residues of this domain (Arg^511^), point to the relevance of the Rab-GAP activity for TBCK function, even though its physiological targets have not been identified.^19^ Indeed, the “Arg finger” is conserved in virtually all TBC domains with GAP activity,^21,22,25^ and it is largely accepted that unconventional TBC domains lacking this residue do not stimulate GTP hydrolysis in Rab proteins but have different functions.^21,22,25^ Of note, one of the human unconventional TBC proteins (TBC1D26) and its *Chlamydomonas reinhardtii* orthologue have a histidine residue in place of the “catalytic” arginine, and no GAP activity was detected in the algal protein.^31^ Mutation of the “catalytic” Arg in TBC domain-containing proteins is commonly used as a tool to investigate their function and identify their substrate Rabs,^22,32–35^ since substitution of this amino acid greatly reduces the catalytic efficiency of these GAPs.^14^

TBCK’s role in controlling cell proliferation/growth and actin cytoskeleton organization has been reported to be mediated by modulation of the mTOR signaling network and transcriptional regulation of components of the mTOR complex.^19^ The mTOR pathway is a growth-regulating network, which in an activated state promotes angiogenesis, cell growth, and cell proliferation. A number of developmental brain disorders, collectively termed “TORopathies”, have been shown to result from dysregulated mTOR signaling.^36,37^ These disorders are characterized by disorganized cortical lamination, seizures and cytomegaly.^38^ Tuberous sclerosis complex (TSC) [MIM PS191100] is caused by heterozygous loss of function mutations in *TSC1* [MIM 605284] or *TSC2* [MIM 191092], encoding proteins that negatively regulate the mTOR pathway. Similarly, germline or postzygotic activating *de novo* mutations in components of the PI3K-AKT3-mTOR pathway cause the overlapping hemimegalencephaly phenotypes megalencephaly-capillary malformation-polymicrogyria [MIM 602501] (usually caused by somatic mutations in *PIK3CA* [MIM 171834]) and megalencephaly-polymicrogyria-polydactyly-hydrocephalus [MIM PS603387] (usually caused by germline mutations in *PIK3R2* [MIM 603157], *CCND2* [MIM 123833, and *AKT3* [MIM 611223]), in addition to their well-documented role in cancer. More recently, germline gain-of-function mutations in *MTOR* [MIM 601231] have been described in two families with features overlapping the megalencephaly and RASopathy spectrum of disorders.^39,40^ These TORopathies are all caused by activation of the mTOR pathway and generally characterized by brain overgrowth at a global (hemimegalencephaly spectrum) or focal level (TSC). In contrast, loss of *TBCK* function is associated with loss of brain volume. In combination with available biochemical and functional data supporting a positive modulatory role of TBCK on mTOR signaling,^19^ our findings suggest that loss of TBCK function has an opposite effect on this signaling network compared to what is observed in other mTOR-related disorders, and importantly, that over-inhibition of the mTOR pathway may lead to this distinct and severe phenotype. Some of the affected individuals were found to have moderate mitochondrial dysfunction which did not meet Walker diagnostic criteria^41^ for a primary respiratory chain disorder and was not associated with increased brain lactate in the three individuals evaluated by magnetic resonance spectroscopy (Table S3). Given that the mTOR pathway is known to positively affect mitochondrial biogenesis^42^ and negatively regulate autophagy/mitophagy (reviewed in Bockaert and Marin 2015^43^) and that several individuals have decreased mitochondrial enzyme levels, we hypothesize that the progression of the disease may be due, in part, to decreased mitochondrial biogenesis and/or increased mitophagy.

In summary, we establish that biallelic mutations in *TBCK* cause a severe neurodevelopmental disorder whose major features include profound developmental delay/cognitive deficit, brain atrophy without microcephaly, dysgenesis of the corpus callosum, white matter signal abnormality, cerebellar vermis hypoplasia, seizures, diminished respiratory function, and distinctive facies. The recent report of a similar individual exhibiting severe developmental disability, poor feeding, abnormal eye movements, epilepsy, facial dysmorphism, severe hypotonia, and diffuse brain atrophy associated with a homozygous canonical splice-site mutation (NM_001163435.1 c.1897+1G>A) in *TBCK^44^* further strengthens this view. Structural and molecular modeling analyses predict a key role for the substituted residue in mediating the Rab-GAP activity of the protein, suggesting that loss of TBCK GAP function is sufficient to cause disease. Of note, many encephalopathies are caused by *de novo* dominant mutations and therefore have low risk of recurrence, so distinguishing TBCK-related encephalopathy is essential for accurate recurrence risk counseling and reproductive planning. Recognition of this disorder will also make it possible to delineate the natural history of TBCK-related disease, and provide more precise prognostic information that is essential to guide decisions about invasive treatments such as tracheostomy and gastrostomy in neonates and young children.

More than three years passed between the identification of *TBCK* as the strongest candidate gene in Family A and the identification of *TBCK* mutations in Families B, C, and D. Importantly, even though two families were sequenced through the same center (University of Washington Center for Mendelian Genomics), their shared phenotype was not initially recognized, in part because the clinicians rightfully focused on different aspects of each affected individual’s condition. The identification of a shared candidate gene by searching Geno_2_MP prompted comparison of the clinical findings and led to delineation of an overlapping phenotype. It is anticipated that matchmaking platforms in development, such as Matchmaker Exchange,^45^ will help to speed up this process, but only if investigators and clinicians worldwide all participate in such data sharing efforts. Historically, ascertaining persons with a highly similar phenotype and then conducting subsequent gene discovery within that group has been a successful method for understanding new Mendelian phenotypes; however, this approach requires the Mendelian phenotype to be common enough that a single investigator could be expected to encounter multiple affected individuals. The small number of individuals with biallelic mutations in *TBCK* identified thus far, and the international assemblage of investigators who identified these individuals, underscores the fact that many, if not most, Mendelian phenotypes now being studied will be more rapidly and successfully tackled by widespread sharing of phenotypes, genotypes, and candidate genes.^13,45,46^

## Supplemental Data

Supplemental data consists of 3 tables.

## Acknowledgments

We thank the families for their participation and support. Our work was supported in part by grants from the National Institutes of Health: National Human Genome Research Institute and the National Heart, Lung and Blood Institute (1U54HG006493 to M.B., D.N., and J.S.; 1RC2HG005608 to M.B., D.N., and J.S.), the National Institute of Neurological Diseases and Stroke (K12NS049453-09 to X.O.G.), the Eunice Kennedy Shriver National Institute of Child Health and Human Development (U54HD083091, Genetics Core and Sub-project 6849 to D.D.), Telethon-Italy (GGP13107 to M.T.) and the Ospedale Pediatrico Bambino Gesù (GeneRare to M.T.). The authors would like to thank the University of Washington Center for Mendelian Genomics and all contributors to Geno_2_MP for use of data included in Geno_2_MP. The authors would also like to thank the Exome Aggregation Consortium and the groups that provided exome variant data for comparison. A full list of contributing groups can be found at http://exac.broadinstitute.org/about.

## Web Resources

The URLs for data presented herein are as follows:

Exome Variant Server (NHLBI Exome Sequencing Project ESP6500): http://evs.gs.washington.edu/EVS/.

Exome Aggregation Consortium (ExAC), Cambridge, MA: http://exac.broadinstitute.org (accessed April 2015).

Geno_2_MP, NHGRI/NHLBI University of Washington-Center for Mendelian Genomics (UW-CMG), Seattle, WA: http://geno2mp.gs.washington.edu (accessed April 2015)

GeneDistiller 2014: http://www.genedistiller.org/

Online Mendelian Inheritance in Man: http://www.omim.org

